# D-serine induces distinct transcriptomes in diverse *Escherichia coli* pathotypes

**DOI:** 10.1101/2021.05.12.443952

**Authors:** James P. R. Connolly, Natasha C. A. Turner, Jennifer C. Hallam, Patricia T. Rimbi, Tom Flett, Mhairi J. McCormack, Andrew J. Roe, Nicky O’Boyle

**Author notes:** These authors contributed equally to this work (James P. R. Connolly, Natasha C. A. Turner, Jennifer C. Hallam).

## Abstract

Appropriate interpretation of environmental signals facilitates niche specificity in pathogenic bacteria. However, the responses of niche-specific pathogens to common host signals are poorly understood. D-serine (D-ser) is a toxic metabolite present in highly variable concentrations at different colonisation sites within the human host that we previously found is capable of inducing changes in gene expression. In this study, we made the striking observation that the global transcriptional response of three *Escherichia coli* pathotypes - enterohaemorrhagic *E. coli* (EHEC), uropathogenic *E. coli* (UPEC) and neonatal meningitis associated *E. coli* (NMEC) - to D-ser was highly distinct. In fact, we identified no single differentially expressed gene common to all three strains. We observed the induction of ribosome-associated genes in extraintestinal pathogens UPEC and NMEC only, and the induction of purine metabolism genes in gut-restricted EHEC and UPEC indicating distinct transcriptional responses to a common signal. UPEC and NMEC encode *dsdCXA* – a genetic locus required for the detoxification and hence normal growth in the presence of D-ser. Specific transcriptional responses were induced in strains accumulating D-ser (WT EHEC and UPEC/NMEC mutants lacking the D-ser-responsive transcriptional activator DsdC), corroborating the notion that D-ser is an unfavourable metabolite if not metabolized. Importantly, many of the UPEC-associated transcriptome alterations correlate with published data on the urinary transcriptome, supporting the hypothesis that D-ser sensing forms a key part of urinary niche adaptation in this pathotype. Collectively, our results demonstrate distinct pleiotropic responses to a common metabolite in diverse *E. coli* pathotypes, with important implications for niche selectivity.

**Importance:** The pathogenic *Escherichia coli* comprise a group of highly specialized bacteria, some of which are capable of disseminating from the intestine and causing disease at other sites within the human host. Chemicals (metabolites) derived from the host and other microorganisms shape the behaviour of *E. coli* in different environments. Here we investigate the changes in gene expression that occur in *E. coli* strains capable (UPEC and NMEC) and incapable (EHEC) of metabolizing D-ser – a metabolite specifically enriched in the urine and in regions of the brain. We show that EHEC, UPEC and NMEC – distinct pathotypes associated with disease in the gut, bladder and brain, respectively, respond in a distinct manner to D-ser. Many of the genes affected by D-ser have been shown to be important during disease, highlighting the importance of varied responses to this common signal in host infection.

## Observation

Pathogenic *Escherichia coli* comprise a diverse, ecologically specialized group of microorganisms capable of causing disease within the intestine – their primary site of colonization – but also at extraintestinal sites such as the brain and bladder. Our previous work described the incompatibility of enterohaemorrhagic *E. coli* (EHEC) with environments rich in the host metabolite D-ser (1). Typically, EHEC cannot metabolize D-ser due to an evolutionary loss of *dsdC*, encoding the D-ser responsive transcriptional activator (2). In addition to growth inhibition by D-ser, the expression of EHECs primary colonisation apparatus, the locus of enterocyte effacement (LEE)-encoded type three secretion system is repressed. It has been reported that D-ser is concentrated in the hippocampus and frontal cortex of the brain (3, 4), while concentrations of D-ser in human urine can reach in excess of 1 mM (5). The toxicity of D-ser to EHEC at such concentrations is hypothesised to be a key factor in restricting EHEC to its preferred gut niche, where D-ser concentrations are extremely low. Indeed, carriage of an intact *dsdCXA* locus for D-ser metabolism in EHEC is extremely rare (1).

The role of D-ser in modulating transcription in EHEC has been extensively investigated, however there are often important intraspecies distinctions in responses to the same metabolite with implications within the context of infection (6–8). Here we describe the transcriptional response of uropathogenic and neonatal meningitis-associated *E. coli* (UPEC and NMEC) to D-ser. Unlike EHEC, these pathotypes typically encode *dsdCXA*, with NMEC strains often carrying two copies of the locus (2). This is believed to be important in pathogenesis in the bladder and brain where D-ser concentrations are higher than that of the gut. We previously described how exposure of EHEC to D-ser results in a global transcriptional shift affecting virulence and inducing stress (1, 9). We therefore hypothesised that D-ser could promote distinct transcriptomes in pathotypes not susceptible to D-ser toxicity. EHEC, UPEC and NMEC were found to display markedly different growth (strong repression of EHEC while no detrimental effects were observed with UPEC or NMEC) rates in M9 minimal medium containing 1mM D-ser, in line with previous reports (Fig. 1A-C). In order to assess the transcriptional response of an actively growing population to D-ser, we opted to spike 1 mM D-ser into the media after 3 hours growth and sample cells after 2 hours of exposure using RNA-seq analysis. Intriguingly, this led to a transient drop in the UPEC growth rate but not NMEC (Fig. 1E and F), possibly as a result of differential metabolic adaptation to the use of D-ser as a carbon source. This surprising common growth inhibition phenotype observed in EHEC and UPEC was reflected by similar numbers of differentially expressed genes (DEGs) (162 and 140 respectively; Fig. 1D-G; Dataset S1). Contrastingly, NMEC growth was not significantly affected and only 55 D-ser-induced DEGs were identified. DEGs belonged predominantly to transport and metabolism/biosynthesis functional categories (Fig. S1). Surprisingly, UPEC and NMEC shared fewer common DEGs (12) than UPEC and EHEC, despite their shared ability to metabolize D-ser (14; Fig. 1G). Importantly, there were no DEGs in common between all three strains, highlighting the individuality in pathotype responses to this metabolite.

**Fig. 1.**
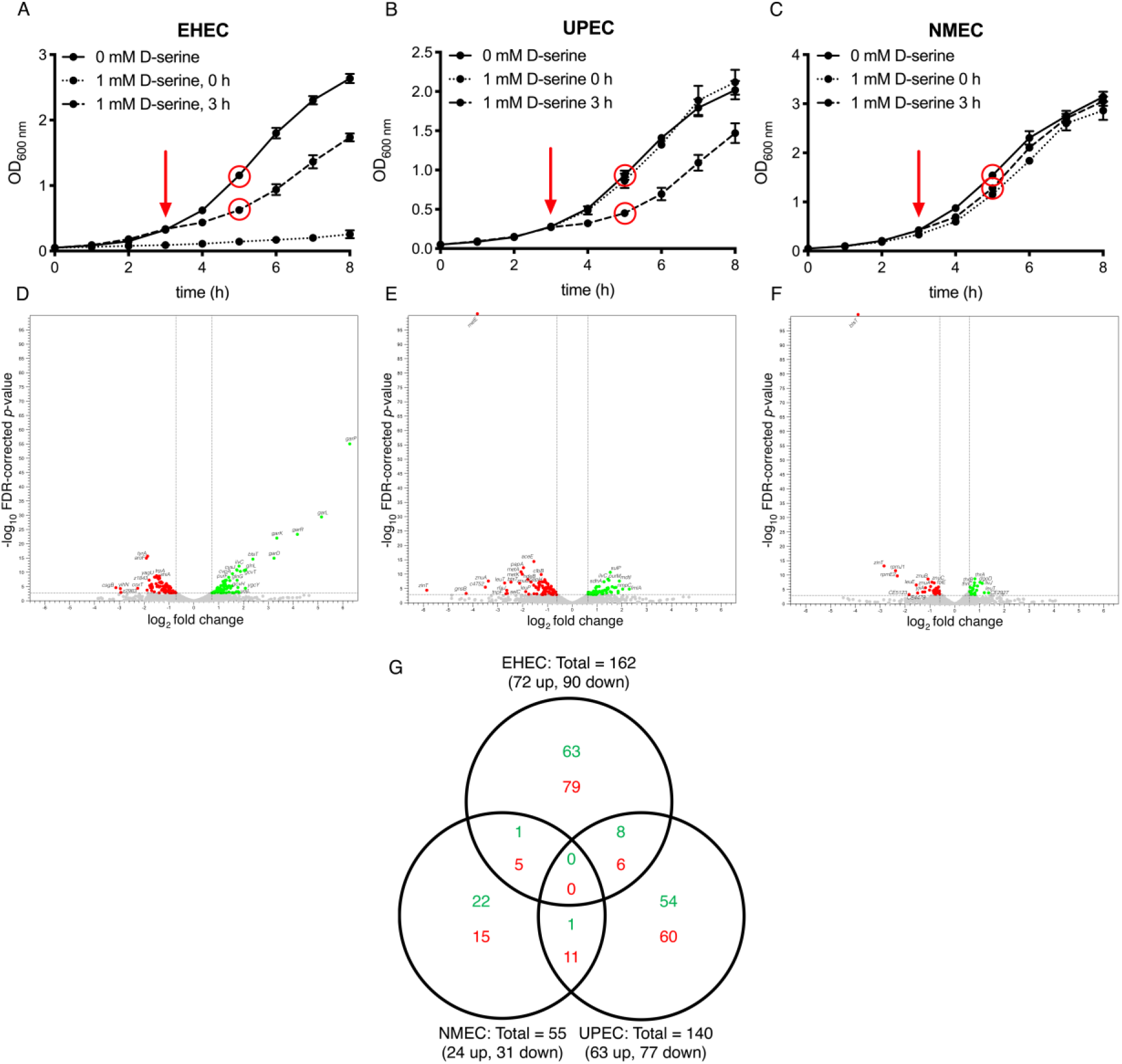
Distinct growth and transcriptional responses of *E. coli* pathotypes to D-serine. (A-C) Growth curves of EHEC, UPEC and NMEC in the presence and absence of 1 mM D-ser, added at 0 h or 3 h (red arrow) post-inoculation as indicated. Red circles indicate the timepoints and samples used for RNA-seq analysis. (D-F) Volcano plots depicting differential gene expression in EHEC, UPEC and NMEC, respectively, following 2 h exposure to 1 mM D-ser. Dots were manually added beyond the axis limits where FDR-corrected *p*-value of zero was obtained. (G) Venn diagram illustrating the distinction between genes differentially expressed following exposure to D-ser in three *E. coli* pathotypes.

The most significantly affected functional categories in all three pathotypes were metabolism/biosynthesis, followed by transport (Fig. S1 and Dataset S2), however the composition of these categories varied significantly between strains. This was highlighted by the fact that the genes/gene clusters bearing the highest level of D-ser-dependent induction were unique to each strain. In EHEC, transport/metabolism of galactarate/glucarate (*gar*) was highly upregulated (Fig. 1D), whereas in UPEC the fimbrial gene *fmlA* – the major subunit of the virulence-associated F9 pilus (10) – was most induced (Fig. 1E) and in NMEC threonine biosynthesis (*thr*) was strongly activated (Fig. 1F). The most downregulated genes in EHEC were associated with curli synthesis (*csg*) and tryptophan biosynthesis (*trp*) (Fig. 1D), while in UPEC and NMEC, a zinc chelator encoded by *zinT*, and zinc transporter *znuABC* were strongly reduced in expression (Fig. 1 E and F). Several genes involved in purine metabolism were upregulated in both EHEC and UPEC, while all three strains exhibited repression of distinct acid tolerance genes. These included *cfa, kgtP*, and *phoH* in EHEC, *gadX* in NMEC, *hdeABD, gadAB, ybaST* in UPEC, *yagU* in both EHEC and NMEC, and *gadC* in both EHEC and UPEC. We predicted that many conserved genes would be inversely regulated in EHEC compared with UPEC/NMEC based on D-ser deamination capability, however only one DEG followed such a pattern – a CstA family pyruvate transporter encoded by *yjiY/btsT* that was upregulated in EHEC and repressed in UPEC and NMEC (Fig. 1E).

In addition to *fmlA*, several DEGs with characterized roles in pathogenesis were affected by D-ser. In UPEC, genes belonging to both operons encoding the *pap* pilus – a type 1 fimbria that is overrepresented in pyelonephritis isolates and functions by binding the globoside glycolipid receptor in the kidney (11) - were repressed. We also observed a striking upshift in ribosomal protein/RNA gene expression in UPEC and NMEC but not in EHEC. Several 50S (*rpl*) and 30S (*rps*) ribosomal protein genes were upregulated in both pathotypes in response to D-ser (Fig. 1E and F). Importantly, many UPEC responses (including upregulation of ribosomal genes, repression of *pap* pilus and activation of cold shock genes) to D-ser have also been reported in response to growth in human urine (12) or during mouse/human urinary tract infection (12–14). Upregulation of ribosomal genes is believed to play a role in disease by facilitating rapid growth via increased translation (14). Thus, these findings support our hypothesis that D-ser sensing is crucial for recognition of the urinary niche by UPEC.

In order to distinguish D-ser transcriptional responses associated with accumulation (analogous to the natural scenario of EHEC) from those driven by metabolism of D-ser, we performed a parallel transcriptome analysis using UPEC and NMEC strains deleted for *dsdC* (the D-ser responsive transcriptional activator gene required for D-ser metabolism(15)). Note, as mentioned above NMEC encodes two copies of *dsdC*, which were both deleted. As predicted, UPEC Δ*dsdC* and NMEC Δ*dsdC1/C2* displayed severe growth arrest in the presence of D-ser (Fig. 2A-C). Exposure resulted in 1345 DEGs in UPEC Δ*dsdC*, 24% of its gene content. Contrastingly, NMEC Δ*dsdC1/C2* displayed 357 DEGs in response to D-ser. The large-scale shift in gene expression in these mutants compared with WT EHEC (162 DEGs) reflects their observed incompatibility with D-ser accumulation (Fig. 2A and B). This is consistent with the observation that carriage of intact *dsdCXA* is extremely common in these extra-intestinal isolates and that they can inhabit environments abundant in D-ser, where detoxification is a prerequisite for success. Accordingly, UPEC Δ*dsdC* and NMEC Δ*dsdC1/C2* displayed induction of greater numbers of stress response genes than their wild type counterparts upon D-ser exposure and accumulation. While some overlap was observed in the responses of WT and *dsdC* mutants to D-ser (including ribosome associated genes – suggesting a sensing mechanism independent of accumulation), the response of the mutants was largely distinct (Fig. 2E). Comparison of *dsdC*-negative UPEC and NMEC mutant response with EHEC identified a core set of 30 D-ser accumulation-dependent DEGs (Fig. 2F). These include induction of galactarate transport and metabolism genes (*gar*), glycine cleavage system genes (*gcv*) (Fig. 2C and D; Dataset S1). This subset of DEGs likely contributes specifically to the growth arrest phenotype associated with D-ser accumulation and is currently under investigation.

**Fig. 2.**
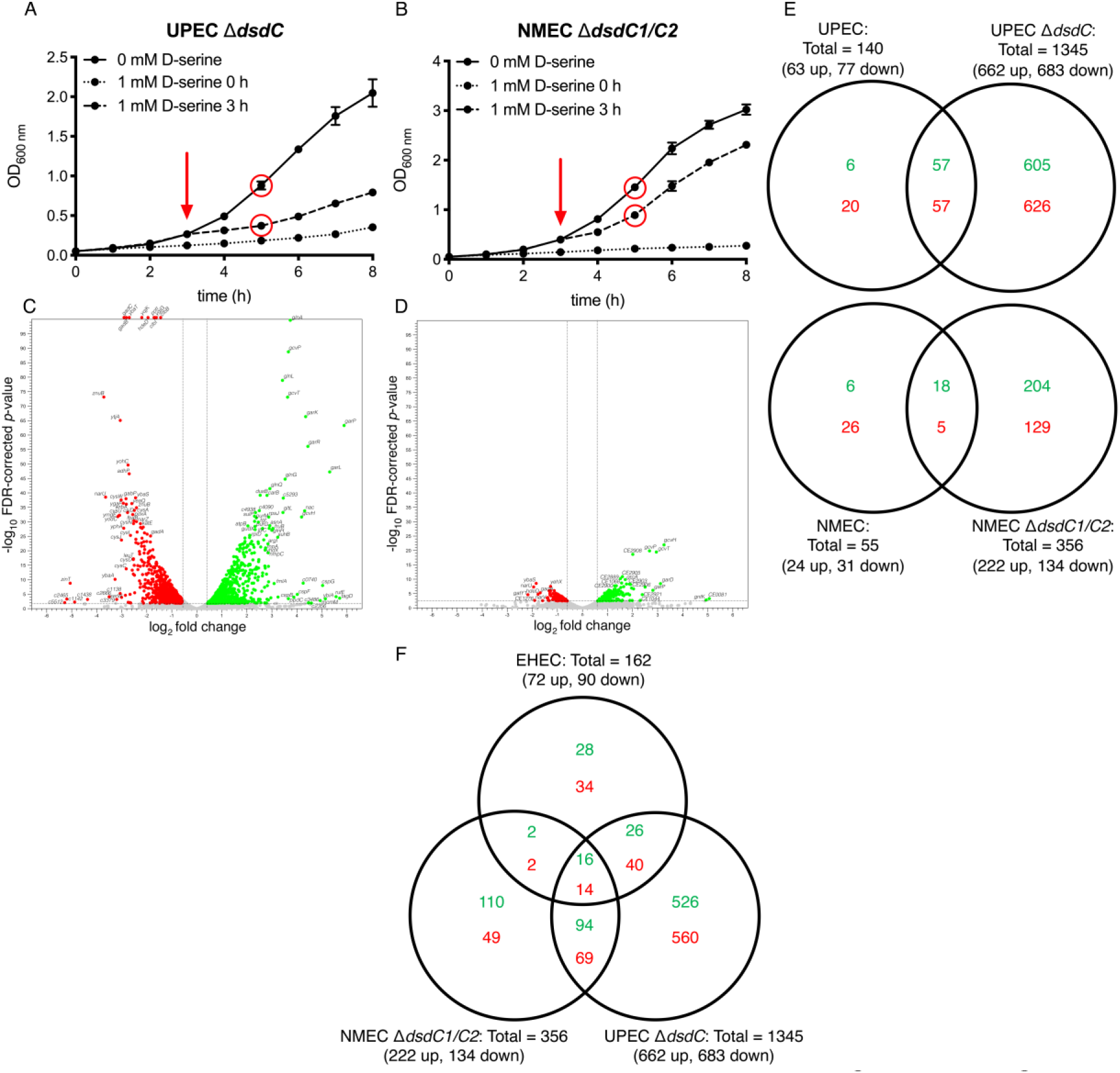
Accumulation of D-ser has variable effects on UPEC and NMEC. (A and B) Growth curves of UPEC Δ*dsdC* and NMEC Δ*dsdC1/C2* in the presence and absence of 1 mM D-ser, added at 0 h or 3 h (red arrow) post-inoculation as indicated. Red circles indicate the timepoints and samples used for RNA-seq analysis. (C and D) Volcano plots depicting differential gene expression following 2 h exposure to 1 mM D-ser. Dots were manually added beyond the axis limits where FDR-corrected *p*-value of zero was obtained. (E) Venn diagram illustrating comparing the effects of D-ser exposure in parental and DsdC-negative deletion mutants of UPEC and NMEC. (F) Venn diagram comparing genes differentially expressed after D-ser exposure in strains lacking DsdC.

## Conclusion

Exposure to D-ser resulted in distinct transcriptional responses in EHEC, UPEC and NMEC, highlighting the diverse strategies employed by different pathotypes in responding to a relevant environmental signal. Large scale subversion of global transcription occurred upon accumulation of D-ser, highlighting the strict requirement for detoxification in strains that occupy niches rich in this metabolite. Importantly, the data correlate with observations made from human studies of the UPEC transcriptome during UTI, suggesting specific responses to D-ser as being highly relevant in a true physiological setting. These surprising nuances in transcriptional responses displayed by distinct but closely related pathotypes highlight the challenges faced when using single prototypic strains in drawing species-level conclusions.

## Acknowledgements

NO’B is supported by a Tenovus Scotland award. JPRC is supported by a Springboard award from the Academy of Medical Sciences [SBF005\1029]. AJR is supported by the Biotechnology and Biological Sciences Research Council [BB/M029646/1, BB/R006539/1].

## Figures

**Fig. S1.**
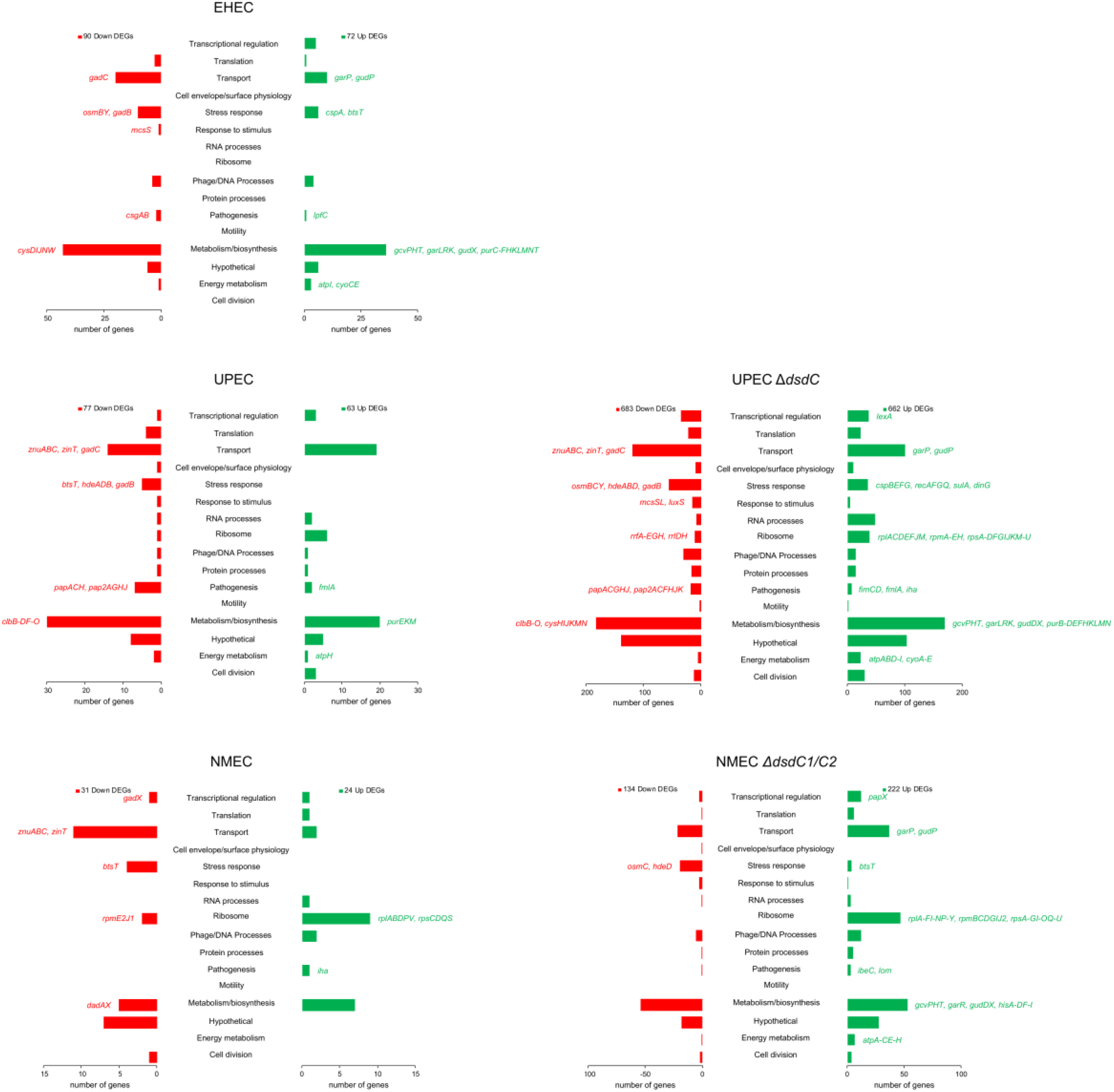
Functional category analysis of differential gene expression in response to D-serine.

